# Actin cytoskeleton self-organization in single epithelial cells and fibroblasts under isotropic confinement

**DOI:** 10.1101/318022

**Authors:** Salma Jalal, Ruby Yun-Ju Huang, Virgile Viasnoff, Yee Han Tee, Alexander Bershadsky

## Abstract

We systematically investigated the principles of actin cytoskeleton self-organization in two cell types, fibroblasts and epitheliocytes, by confining isolated cells on isotropic adhesive islands of varying size. In fibroblasts, we previously described that an initially circular pattern of circumferential actin dynamically evolves into a radial pattern of actin bundles that spontaneously transforms into a chiral pattern, before finally producing parallel linear stress fibres. We now show that progression from circular to chiral actin patterns depends on cell projected area and rarely occurs on small islands. Epitheliocytes however, did not exhibit succession through all the actin patterns described above even on large islands. Upon confinement, the actin cytoskeleton in non-keratinocyte epitheliocytes is arrested at the circular stage, while in keratinocytes it can progress as far as the radial pattern but still cannot break symmetry. Epithelial-mesenchymal transition pushed actin cytoskeleton development from circular towards radial patterns but remains insufficient to cause chirality. Surprisingly, small doses of G-actin sequestering drug, latrunculin A, induced chiral swirling in keratinocytes. During this swirling, keratin filaments follow actin and also demonstrate chiral swirling movement. Elimination of the keratin network by genetic silencing of Type II keratins, however, did not affect the self-organization of the actin cytoskeleton.

## Introduction

Differences in the morphogenetic behaviour between fibroblasts and epithelial cells have been well documented. *In vivo*, epithelial cells possess apico-basal polarity and are organized into sheets of cells that are connected to each other via E-cadherin mediated cell-cell junctions and attached to the basement membrane via integrin mediated adhesions. In contrast, fibroblasts are embedded and attached to the extracellular matrix of connective tissue and do not readily adhere to neighbouring cells. The difference between morphology and migratory characteristics of epithelial cells and fibroblasts are accompanied by differences in their actin cytoskeleton organization. The most prominent element of the actin cytoskeleton in epithelial cells is the adhesion belt associated with the cell-cell junctions and located at the apical zone of the cells ^1-3^. In typical fibroblasts the actin cytoskeleton is organized into parallel linear actomyosin bundles (stress fibres) associated with cell-matrix adhesions (focal adhesions) ^4-6^. Consistently, upon epithelial-mesenchymal transition (EMT) the circumferential bundles in epithelial cells are succeeded by a system of parallel stress fibres ^7,8^. Several actin regulatory proteins have been shown to be involved in EMT ^9-11^.

It is well documented that outside-in signalling from adhesion receptors directs the formation of the actin cytoskeleton within cells ^2, 12^. Therefore, it is possible that differences in actin cytoskeleton self-organization between epithelial cells and fibroblasts are determined by the differences in available extracellular adhesions (cell-cell and cell-matrix versus cell-matrix only). On the other hand, it is also possible that the actin cytoskeleton itself has intrinsically different morphogenetic potentials in these two cell types. To reveal the difference in the morphogenetic potential of the actin cytoskeleton independently of differences in adhesion conditions, we study the self-organization of the actin cytoskeleton in a standardized microenvironment provided by microfabricated adhesive patterns ^13, 14^. Under such conditions, cells are deprived of adhesions to neighbouring cells and only contact extracellular matrix proteins (specifically fibronectin or collagen). To dissect the processes of actin cytoskeleton self-organization from potential interplay with cell shape changes during migration, we confined cells to isotropic circular adhesive islands so that they acquire discoid morphology and do not change shape.

We have previously used this system of isotropic confinement to investigate the process of actin cytoskeleton self-organization that leads to formation of parallel linear stress fibres in isolated fibroblasts ^15^. This setting revealed unique patterns of actin development with initially symmetric *circular* patterns consisting of circumferential bundles, that progressed into *radial* patterns composed of bundles of actin anchored at focal adhesions that grow radially inwards from the cell periphery. These patterns were followed by a symmetry breaking process that produced a novel *chiral* pattern in which all radial bundles of actin tilted in one direction. This tilting converts the centripetal actin flow into a swirling motion that persists until arrays of parallel stress fibres are formed (the *linear* pattern).

A number of studies have found that geometry and size of adhesive islands determine a variety of different aspects of cellular organization and function ^16-20^. Therefore, in search of external factors that may determine the self-organization of the actin cytoskeleton we first investigated the effect of a basic scaling parameter, cell projected area. We have indeed found that progression through the successive actin patterns described in fibroblasts depends on cell projected area and correlate with focal adhesion formation. We further investigated systematic differences in actin cytoskeleton self-organization between epithelial cells and fibroblasts as well as between different types of epithelial cells. In particular we found that the morphogenetic repertoire of actin cytoskeleton patterns in epithelial cells differs from that of fibroblast, and that EMT can push development of the actin cytoskeleton in epithelial cells towards a fibroblast-type pathway.

The presence of the cytokeratin network in epithelial cells, its depletion during EMT and absence in fibroblasts, makes its interplay with the actin cytoskeleton a possible candidate to explain the differences in the actin cytoskeleton development between the two cell types. However, we demonstrated that the pattern of actin cytoskeleton self-organization in keratinocytes does not require cytokeratin filaments.

## Results

### Actin cytoskeleton self-organization in isolated fibroblasts depends on projected cell area

In isolated human foreskin fibroblasts (HFF) confined on circular adhesive islands of 1800μm^2^ in area covered with fibronectin, the actin cytoskeleton initially self-organizes into a *circular* pattern of circumferential actin bundles with some lamellipodial and filopodial protrusions (Figure 1A). Sometimes these cells demonstrate sparse ventral stress fibres that are not organized into arrays (Movie 1). Within a few hours this circular pattern is succeeded by a symmetric *radial* pattern formed by radial fibres (RFs) that grow inwards from focal adhesions and transverse fibres (TFs) oriented orthogonally to the RFs and parallel to the cell edge (Figure 1A). After some time a symmetry breaking event occurs and all RFs tilt in the counter-clockwise direction forming the *chiral* pattern, before actin bundles eventually organize into the *linear* pattern containing parallel linear stress fibres (Figure 1A) similar to those seen in fibroblasts polarized on planar fibronectin coated substrates (Figure S1A). This development and progression through circular to linear actin patterns is consistent with previously published data ^15^.

**Figure 1:**
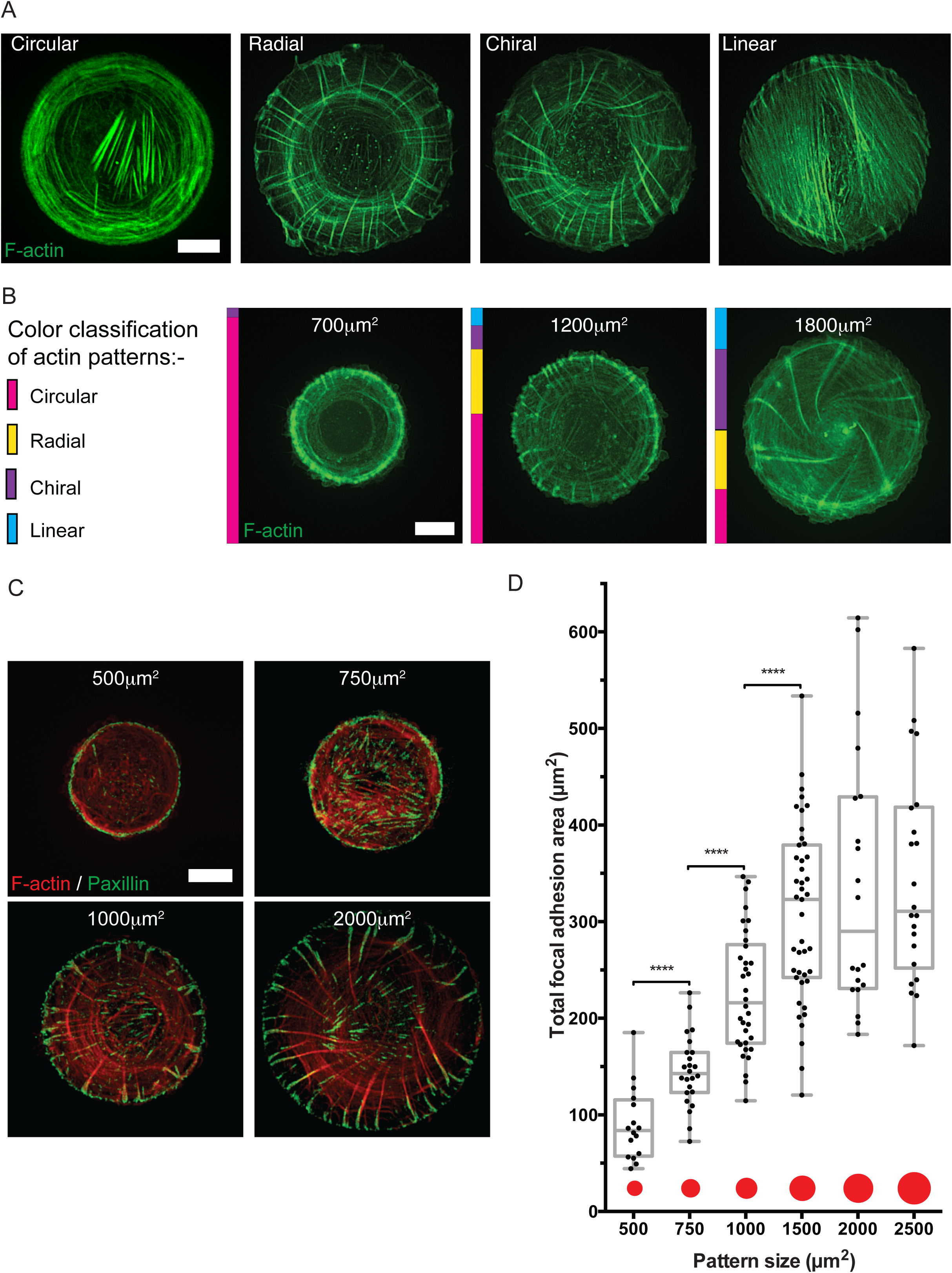
Evolution of actin cytoskeleton self-organization in fibroblasts confined on adhesive islands of varying size. **A)** Representative images showing F-actin (phalloidin staining) distribution in HFF fixed 6hrs after seeding on fibronectin coated islands of 1800μm2. Cells normally initially organize a *circular* pattern consisting of circumferential actin bundles. After ~3hrs the *circular* pattern is succeeded by the *radial* actin pattern upon radially symmetric growth of actin bundles (radial fibers, RFs) inwards from the peripheral focal adhesion. Simultaneously, circumferential actin bundles parallel to the cell edge (transverse fibers, TFs) move centripetally along RFs. After ~6hrs in a majority of cells, all radial fibers spontaneously tilt in one direction to form the *chiral* actin pattern, which persists until actin stress fibers ultimately linearize in the *linear* actin pattern. See Movie 1. Scale bar, 10μm. **B)** Representative frames showing actin pattern development in HFF expressing Lifeact-GFP seeded on 700, 1200 or 1800μm^2^ fibronectin coated islands and imaged overnight. Fractions of cells with actin cytoskeleton demonstrating circular, radial, chiral or linear pattern by the end of filming (8-12hrs) on each island size are represented by the coloured bars. 25, 40 and 100 cells for the 700, 1200 and 1800μm^2^ island sizes respectively were filmed from which actin patterns were scored. Scale bar, 10μm. **C)** Representative images showing focal adhesion (paxillin immunostaining) and F-actin (phalloidin staining) distribution in fibroblasts fixed 8hrs after seeding on fibronectin coated islands of different sizes. Scale bar, 10μm. **D)** Box and whiskers plot showing the total focal adhesion area per cell in cells imaged under conditions of **C**. The total focal adhesion area was measured in 16, 24, 34, 43, 20 and 22 cells on island sizes of 500, 750, 1000, 1500, 2000 and 2500 μm^2^ respectively. Mann-Whitney tests were used to assess significance between the total focal adhesion areas in pairs of island sizes; statistically significant differences are indicated by asterisks corresponding to *p* values as follows: * *p*<0.05, ** *p*<0.01, *** *p*<0.001, **** *p*<0.0001.

Given that different cell types have differential capacities to spread on planar adhesive surfaces (coated with fibronectin or collagen, Figure S1B), adhesive islands of different sizes (generated by either microcontact printing or stencilling, see Materials and Methods and Figure S1C) were used to systematically assess whether cell projected area affects actin cytoskeleton self-organization.

When HFF expressing Lifeact-GFP were confined on small adhesive islands (700μm^2^ in area), actin cytoskeleton development was restricted to circular patterns (represented by magenta bars) in 24 out of 25 cells imaged overnight. On medium sized adhesive islands (1200μm^2^ in area), all described actin patterns could be seen during overnight imaging with the proportion of fibroblasts displaying the radial, chiral and linear patterns represented by yellow, purple and cyan bars respectively. At this island size the circular pattern was still demonstrated by more than 50% of cells but the remaining fraction of cells demonstrated progression to the radial, chiral and linear actin patterns. Finally, on islands of 1800μm^2^ the fraction of cells demonstrating the circular pattern was less than a quarter, while the majority of cells are able to develop the chiral pattern (Figure 1B).

Transition from the circular to radial patterns of actin involves the growth of RFs anchored at focal adhesions. To further probe how spread area affects actin cytoskeleton self-organization we examined its effects on focal adhesions visualized by paxillin antibody staining (Figure 1C). Total focal adhesion area in fibroblasts confined on adhesive islands increased with cell projected area. There was a strong statistically significant increase of total focal adhesion area in cells on patterns with areas of 500, 750, 1000 and 1500μm^2^ (Figure 1D). Of note, the increase in total focal adhesion area was somewhat greater than the increase in projected cell area. In particular, there is a mild statistically significant increase between total focal adhesion area normalized by cell-projected area between patterns with areas of 500 and 1000μm^2^ (Figure S1D). The increase in total focal adhesion area approached a plateau at the island size of 1500μm^2^, so that the total focal adhesion area in cells on islands of larger sizes (2000 and 2500μm^2^) remained constant (Figure 1D), while normalized total focal adhesion area decreased (Figure S1D).

### Non-keratinocyte epithelial cells confined on circular adhesive islands demonstrate only initial stages of actin cytoskeleton self-organization

To examine actin cytoskeleton self-organization in epithelial cells confined on circular adhesive islands, a selection of cell lines from different tissues of origin (including mammary, bladder and skin) was used. Epithelial cell morphology in standard culture is different from fibroblasts, in that cells do not elongate and remain discoid (Figure S1A). Of note, the spreading capacity of these lines (MCF-7, MCF-10A and NBT-II) on planar matrix coated substrates is much lower than that of fibroblasts (Figure S1B). All non-keratinocyte epithelial cells tested, preferentially filled smaller adhesive islands and did not fill larger pattern sizes well (Table S1). In epithelial cells not originating from skin imaged overnight or fixed 20-24 hours after seeding, actin cytoskeleton development was mostly restricted to the circular pattern characterized by circumferential actin bundles and protrusive membrane activity at the cell periphery (Figure 2 and S2). Sparse non-organized ventral stress fibres sometimes observed in these cells (white arrowhead, Figure 2) were usually more dynamic than ventral stress fibres in fibroblasts (Movie 2 compared to Movie 1).

**Figure 2:**
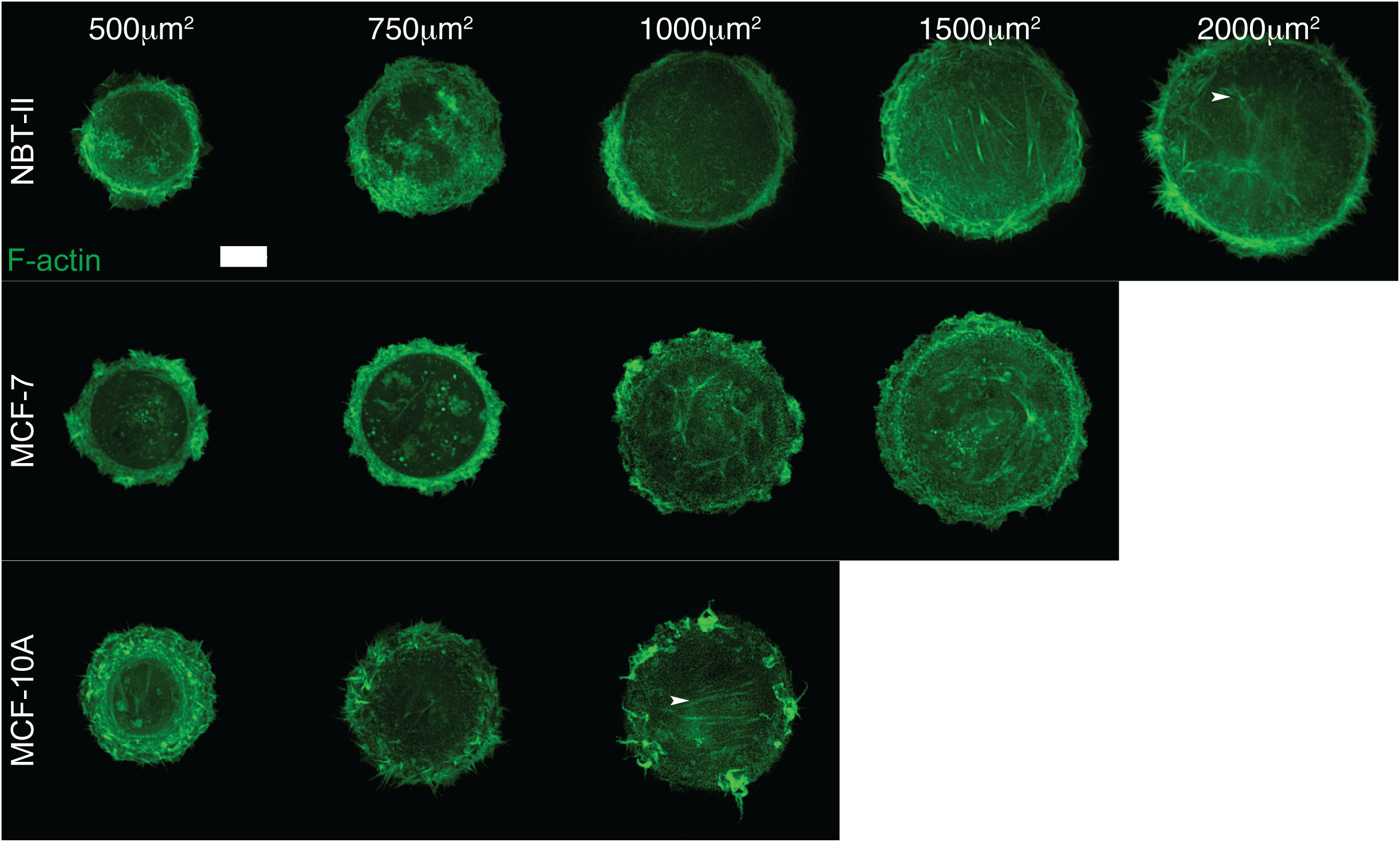
Actin cytoskeleton pattern in non-keratinocyte epithelial cells confined on adhesive islands of varying size. Representative frames of epithelial cells expressing Lifeact-GFP that filled the circular shape when seeded on adhesive islands of different sizes after being imaged overnight. MCF-10A and MCF-7 were seeded on collagen coated substrates while NBT-II were seeded on fibronectin coated substrates. Blank frames correspond to situations when cells did not fill circular islands of the specified size (see Table S1). Scale bar, 10μm.

Next, we investigated how the actin cytoskeleton in epithelial cells changes during EMT and in particular whether the actin pattern of self-organization typical for fibroblasts can be reproduced in epithelial cells undergoing EMT. Epidermal growth factor (EGF) induction of EMT in rat bladder carcinoma cells, NBT-II was used as a model system ^21^. In a colony assay, treatment of EGF induced loosening and partial scattering of NBT-II cell colonies (Figure S3A). In agreement with previous studies, EMT was accompanied by upregulation of Slug and downregulation of E-cadherin (Figure S3B).

Wild type untreated NBT-II cells spread on fibronectin coated islands, mainly demonstrated circumferential bundles of actin and membrane protrusion along with some sparse ventral stress fibres (Figure 3A). After treatment with EGF for at least one day to induce the EMT phenotype, the actin cytoskeleton is able to self-organize into a radial actin pattern (Figures 3A and S3C). This pattern contains RFs originating from focal adhesion-like structures resembling those in fibroblasts (Figure 3A compared to 1A). Unlike the radial actin pattern in fibroblasts, the RFs in EGF treated NBT-II cells are connected to a centrally located dynamic actin array consisting of numerous actin foci connected by thin actin bundles (Figure 3A). This array appeared to assemble at the periphery and disassemble in the central area of the cell thus demonstrating a centripetal flow (Movie 3). Similar to fibroblasts, the self-organization of actin into radial patterns in EGF treated NBT-II cells showed size dependence (Figure 3B). Only 2 out of 22 cells imaged overnight confined on small (500 and 750μm^2^) adhesive islands developed RFs, while 3 out of 15 cells imaged on medium sized islands (1000μm^2^) and 6 out of 16 treated cells imaged overnight on large islands (1500, 2000 and 2500μm^2^) were able to develop RFs (Figure 3B).

**Figure 3:**
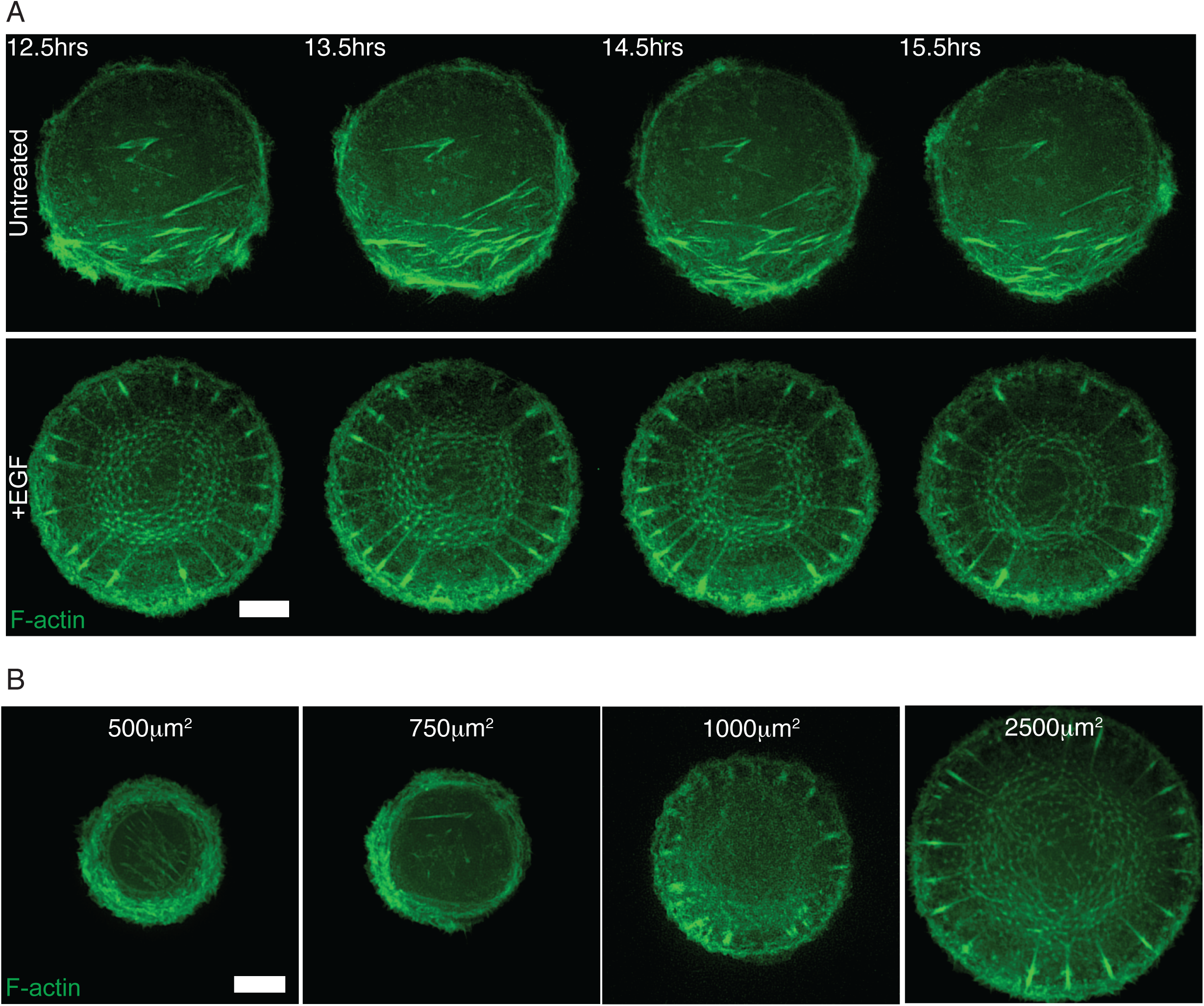
Non-keratinocyte epithelial cells (NBT-II) acquire the ability to form a radial actin pattern after epithelial-mesenchymal transition. **A)** Timelapse series showing actin cytoskeleton in control untreated NBT-II cells expressing Lifeact-GFP and seeded on 1800μm^2^ fibronectin coated islands (upper panel), and NBT-II cells pretreated with 100ng/ml Epidermal Growth Factor (EGF) for 3 days and replated on 1800μm^2^ fibronectin coated islands in the continued presence of EGF (lower panel). Time in hours after seeding is indicated in the upper left corner of frames; the first frame corresponds to the stage when untreated cells filled the entire area of the adhesive circular island. See Movies 2 and 3. Scale bar, 10μm. **B)** Representative frames of actin pattern formation in NBT-II cells replated on fibronectin coated islands of different sizes after 3 days of treatment with 100ng/ml EGF. Cell were transfected with Lifeact-GFP one day before replating and imaging overnight. Scale bar, 10μm.

To further examine the effect of EMT on the morphogenetic potential of the actin cytoskeleton in epithelial cells confined on adhesive islands, we studied a series of ovarian cancer cell lines with different EMT status ^22, 23^. We used four cell lines that differed in their EMT score (Figure S4A). The EMT score is based on transcriptomic analysis of 218 genes (Epithelial: 170, Mesenchymal: 48), and ranges from -1 for the most epithelial phenotype up to +1 for the most mesenchymal phenotype ^22, 23^. Similar to other non-keratinocyte epithelial cell lines tested, the ovarian cancer cell line with epithelial phenotype (PEO1, Figure S4B) only developed circular actin patterns (Figure S4C) and generally did not fill the entire area of large adhesive islands (Table S2). In contrast, HEYA8 cells with the most mesenchymal phenotype (Figure S4B) demonstrated radial or linear actin patterns in the majority of imaged cells (Figures S4C and S4D). The cell lines with intermediate EMT scores (intermediate epithelial, OVCA429 and intermediate mesenchymal, SKOV3) were able to cover the entire surface of large adhesive islands and sometimes formed RFs albeit with lower frequency than the mesenchymal type cell line, HEYA8. On large adhesive islands (1500, 2000, 2500μm^2^), 6 out of 26 imaged OVCA429 and 3 out of 47 imaged SKOV3 were able to develop RFs compared to 29 out of 39 for HEYA8 (where 7 of those cells demonstrated transition to the linear pattern, Figure S4D). Similar to fibroblasts, on small adhesive islands (500, 750, 1000μm^2^) all of these cells were predominantly circular.

### Actin cytoskeleton self-organization in keratinocytes reproduces the radial but not chiral actin patterns

Actin cytoskeleton self-organization in the keratinocyte cell line, HaCaT, a spontaneously immortalized human cell line from skin that is capable of normal terminal differentiation ^24^ was examined. On small adhesive islands there is no clear difference between keratinocytes and fibroblasts since the actin cytoskeleton in both cell types is predominantly circular. As the size of adhesive islands increases so too does the proportion of keratinocytes able to develop RFs in the radial pattern (Figure 4A). The actin cytoskeleton self-organization in keratinocytes only develops as far as the radial pattern and never showed transitions to chiral actin patterns (Figure 4B and Movie 4). The linear actin pattern (final stage of actin cytoskeleton development in fibroblasts) was never observed in HaCaT cells plated and imaged overnight on fibronectin islands. Similar to the situation in EGF treated NBT-II cells, EMT induction in HaCaT cells was insufficient to induce chirality in the actin cytoskeleton (Figure S4). HaCaT cells however, have the potential to develop chiral organization of the actin cytoskeleton. We found that treatment of these cells with a very small dose of actin monomer binding drug, latrunculin A (LatA) was sufficient to trigger the transition from radial to chiral actin patterns (Figure 4C). In keratincoytes filmed overnight on 1800μm^2^ fibronectin islands, 12 out of 40 cells demonstrated chiral actin cytoskeleton swirling. Unexpectedly, the directionality of chiral swirling induced by LatA treatment was clockwise (Figure 4C and Movie 5), opposite to “normal” counter-clockwise swirling characteristic of fibroblasts (Movie 1 and ^15^).

**Figure 4:**
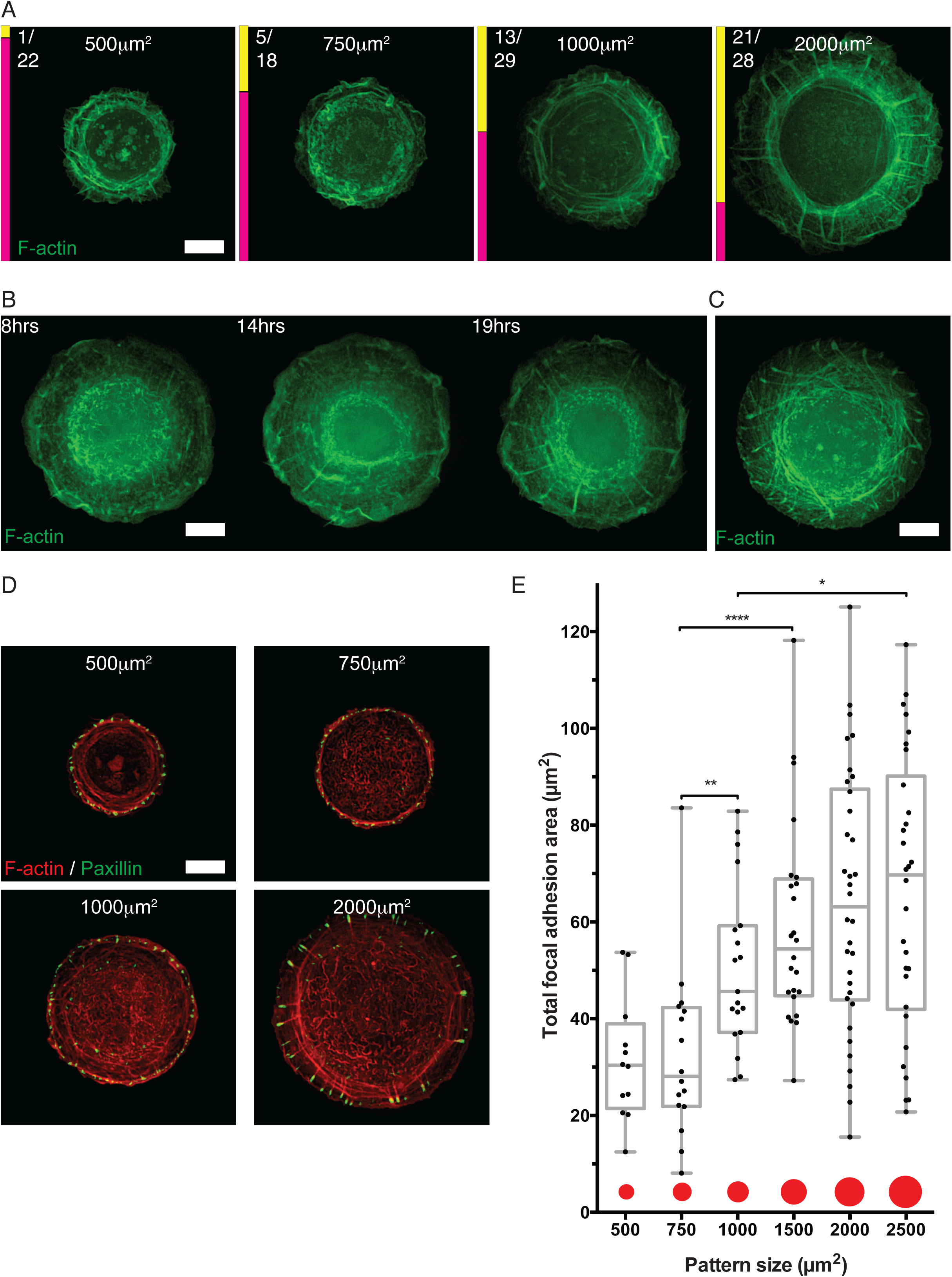
Actin cytoskeleton self-organization in keratinocytes confined on adhesive islands of varying size. **A)** Representative frames showing actin patterns in keratinocytes (HaCaT) expressing Lifeact-GFP or mRuby-Lifeact (pseudo-coloured green) seeded on fibronectin coated islands of different sizes and imaged overnight. The fractions of cells on each island size that demonstrated the circular or radial actin pattern are represented by the magenta or yellow bars respectively. Numbers of scored cells (radial/ total) pooled from 6 independent experiments are shown next to the yellow bars. Scale bar, 10μm. **B)** Timelapse series showing actin cytoskeleton development in HaCaT cells expressing Lifeact-GFP seeded on 1800μm^2^ fibronectin coated islands. Time in hours after seeding is indicated in the upper left corner of frames. See Movie 4. Scale bar, 10μm. **C)** Cells from the same dish as imaged in **B** were treated with 100nM Latrunculin A (LatA) 21hrs after seeding and imaged for an additional 3hrs. Representative frame showing development of the chiral pattern in a fraction of treated cells. See Movie 5. Scale bar, 10μm. **D)** Representative images showing focal adhesion (paxillin immunostaining) and F-actin (phalloidin staining) distribution in HaCaT fixed and stained 24hrs after seeding on fibronectin coated islands of different sizes. Scale bar, 10μm. **E)** Box and whiskers plot showing the total focal adhesion area per cell from cells imaged under conditions of **D**. The total focal adhesion areas were measured in 12, 16, 19, 24, 34 and 30 cells pooled from 2 independent experiments plated on islands of 500, 750, 1000, 1500, 2000 and 2500μm^2^ in area respectively. Mann-Whitney tests were used to assess significance between the total focal adhesion areas in pairs of island sizes; statistically significant differences are indicated by asterisks corresponding to *p* values as follows: * *p*<0.05, ** *p*<0.01, *** *p*<0.001, **** *p*<0.0001.

The total focal adhesion area in HaCaT cells was drastically smaller than in fibroblasts (Figure 4D compared to Figure1C). Analysis of variance (one-way ANOVA with Tukey’s multiple comparison test) revealed significant differences between mean total focal adhesion area of fibroblasts and keratinocytes on all pattern sizes (p<0.0001), except for the smallest pattern area (500μm^2^) where there is no significant difference. However, similar to fibroblasts, keratinocytes confined on circular fibronectin islands also demonstrated dependence of total focal adhesion area on projected area of the cell. There is a statistically significant difference between the total focal adhesion area in two groups of sizes: small (500, 750μm^2^) and relatively large (1000, 1500, 2000, 2500μm^2^) islands (Figure 4E).

### The keratin filament network follows chiral swirling of the actin cytoskeleton but does not affect actin cytoskeleton self-organization

Given that the keratin network is the most abundant and diverse cytoskeletal system in epithelial cells ^25^ and that during the process of EMT endogenous expression of keratin is downregulated in favour of vimentin expression ^22, 23, 26, 27^, the intermediate filament network’s interplay with the actin cytoskeleton was investigated. In keratinocytes plated on 1800μm^2^ fibronectin coated islands the assembly of keratin filaments follows the spatiotemporal pattern previously described in Kolsch et al ^28^. Filaments begin to form near the cell periphery and then move centripetally to be incorporated into a dense network in the central part of the cell (Figure 5A and Movie 6). The chiral swirling of the actin cytoskeleton in keratinocytes induced by mild LatA treatment (Figure 4C) made it possible to investigate the relationship between keratin and actin filament movements. Our observations showed that in keratinocytes demonstrating chiral actin swirling, the keratin cytoskeleton followed the swirling movement of actin (Figure 5B and Movie 7).

**Figure 5:**
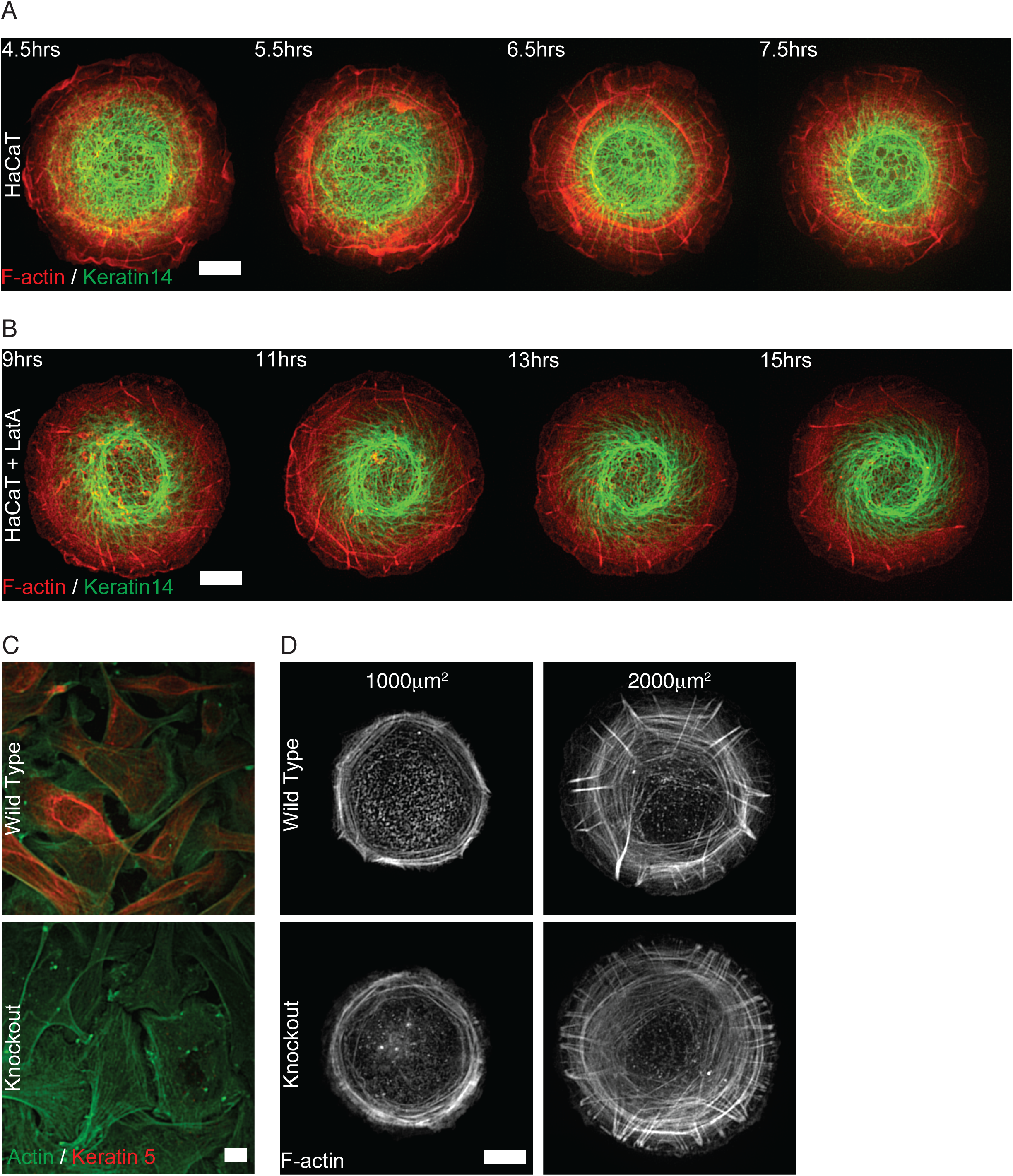
Interplay between cytokeratin and actin cytoskeleton self-organization in keratinocytes confined on adhesive islands. **A)** Timelapse series showing cytoskeleton development in HaCaT cells expressing mEmerald-Keratin14 with mRuby-Lifeact seeded on 1800μm^2^ fibronectin coated islands. Time in hours after seeding is indicated in upper left corner of each frame. Scale bar, 10mm. See Movie 6. Scale bar, 10μm. **B)** Timelapse series showing cytoskeleton progression in Latrunculin A treated HaCaT cells expressing mEmerald-Keratin14 with mRuby-Lifeact seeded on 1800mm^2^ fibronectin coated islands. 20nM of Latrunculin A was added to cells 8hrs after seeding. Time in hours after seeding is indicated in upper left corner of each frame. See Movie 7. Scale bar, 10μm. **C)** Representative images showing Actin (β-actin immunostaining) and Keratin5 (cytokeratin5 immunostaining) in murine keratinocytes with normal (Wild Type) or depleted (Knockout) cytokeratin expression fixed after seeding on collagen coated coverslips. Keratin5 staining was absent in Knockout cells. Scale bar, 10μm. **D)** Representative images showing actin patterns in murine keratinocytes with normal (Wild Type) or depleted (Knockout) cytokeratin expression. Images show F-actin (phalloidin staining) 24hrs after seeding on collagen coated islands of different sizes. Scale bar, 10μm.

In spite of the clear interplay between the actin and keratin cytoskeletons, the removal of the keratin network did not obviously affect actin cytoskeleton self-organization. We compared actin cytoskeleton dynamics in a normal murine keratinocyte cell line (Wild Type, WT) and in a line lacking all keratins (Knockout, KO) (Figure 5C and ^29, 30^). The actin cytoskeleton patterns visualized in both the murine keratinocyte lines 20-24 hours after plating on collagen coated islands was essentially similar to actin cytoskeleton self-organization in the human keratinocyte cell line, HaCaT on fibronectin coated islands (Figure 5D compared to Figure 4A). In both WT and KO keratinocytes plated on small collagen islands (500, 750 and 1000μm^2^) the circular actin pattern dominated in 77 out of 94 and 28 out of 40 cells for WT and KO cells respectively. While in keratinocytes plated on large collagen islands (1500, 2000 and 2500μm^2^) the radial actin pattern developed in 68 out of 73 and 102 out of 122 cells for the WT and KO cells respectively (Figure 5D). We did not observe the development of the chiral actin pattern in any of these cases.

## Discussion

Recent studies of actin cytoskeleton self-organization in fibroblasts confined to circular adhesive islands revealed a rich repertoire of cytoskeletal patterns that can be generated as well as rules underlying the development of these patterns in time ^15^. This approach helped to dissect the intrinsic potential of the actin cytoskeleton for self-organization from the effects that emerge due to cell shape changes and cell migration. So far, only fibroblasts have been investigated under such conditions. In the present study we have analysed the actin cytoskeleton self-organization in a variety of epithelial cell lines under conditions of confinement to circular adhesive islands.

Among the basic parameters that could affect the actin cytoskeleton self-organization is the cell spreading area. Upon spreading under unconstrained conditions the maximal cell spreading area should scale with parameters related to cell size such as cell volume and surface area. Indeed multinucleated cells plated on planar substrates always have larger cell projected areas than their mononucleated counterparts ^31^. The interrelationship between the cell projected area and organization of the actin cytoskeleton, development of traction forces and assembly of focal adhesions was observed in many prior studies ^18-20, 32-34^. Therefore, the optimal development of the actin cytoskeleton requires a sufficient spreading area that could be different for different cell types. In order to properly compare the morphogenetic potential of the actin cytoskeleton in different cell types, it is essential to investigate the process in cells confined to adhesive islands with different areas.

We found that all cells confined to adhesive islands of low cell projected area organized their actin cytoskeleton into a circular actin pattern regardless of type. Differences in actin cytoskeleton development only became apparent in cells confined in large adhesive islands when three distinct types of behaviour were identified in: 1) fibroblasts, 2) keratinocyte epithelial cells and 3) epithelial cells not originating from skin. When plated on sufficiently large adhesive islands fibroblasts are able to efficiently reproduce the full set of actin patterns described previously by progressing to the symmetric radial and then breaking symmetry into the chiral and linear actin patterns ^15^. In contrast, keratinocytes are only able to develop as far as the radial actin pattern while non-keratinocyte epithelial cells cannot even develop past the circular pattern. Epithelial-mesenchymal transition rescued the inability of non-keratinocyte epithelial cells to transition to a radial actin pattern by promoting the development of radial fibres, but remained insufficient to induce development of left-right asymmetry typical of fibroblasts under these conditions.

The main factors participating in self-organization of the actin cytoskeleton are nucleation of polymerization of actin filaments forming radial actin bundles and centripetal flow caused by myosin II driven contractility of transverse circumferential actin bundles. Focal adhesions are the cell organelles that efficiently nucleate the assembly of actin filaments. At the early stages of focal adhesion assembly they are associated with proteins of the Arp2/3 complex ^35-37^, while mature focal adhesions nucleate the associated actin bundles via formin activity ^15, 38-40^. The focal adhesions may not only generate radial fibres but also induce their tilting, which finally leads to the development of chiral swirling. The formins localized to focal adhesions could be responsible for filament twisting coupled with polymerization, which can underlie the mechanism of radial fibre tilting ^15^. Fibroblasts form much larger focal adhesions than epithelial cells, in particular keratinocytes, which could be responsible for the relatively inefficient development of the radial and chiral actin patterns in epithelial cells. At the same time, the keratinocytes have the potential to generate chiral swirling as can be seen in experiments with induction of swirling by low doses of latrunculin A. Even though the direction of latrunculin A induced swirling in keratinocytes is opposite to that of fibroblasts, the resultant actin pattern is qualitatively similar to that of fibroblasts. The molecular mechanisms that are responsible for the differences between the morphogenetic potential of fibroblasts and epithelial cells remain to be determined.

Since keratinocytes differ from fibroblasts by expressing significant amounts of keratin intermediate filament proteins that are organized into well-developed filament networks ^41^, we examined the role of keratins in the development of the actin cytoskeleton. Keratin filaments are physically connected to the actomyosin network which can be inferred from published data ^42, 43^, and our own results showing that keratin filaments follow the actin cytoskeleton swirling in latrunculin A treated cells. Nevertheless, we have found that depletion of keratin proteins ^30^ that resulted in the disappearance of the keratin filament network did not change the actin filament organization in murine keratinocytes. It still remains uncertain whether deficiencies in the development of the actin cytoskeleton of epithelial cells depend instead on the absence of another intermediate filament protein, vimentin. Further work is required to elucidate this.

Our study has revealed clear and reproducible differences between development of the actin cytoskeleton in fibroblasts and epithelial cells. Such differences were evident in a very simple experimental system when cells are deprived of contacts with each other and of the possibility to migrate. Moreover the adhesive islands confining cells were completely isotropic and did not provide any asymmetric cues. Other cytoskeletal elements such as intermediate filaments did not apparently affect the process of actin cytoskeleton self-organization. Thus, such conditions allowed us to determine the most basic differences between the processes of actin cytoskeleton self-organization in different cell types. Besides the apparent differences between fibroblasts and epithelial cells in general, we found that even different types of epithelial cells can be distinguished by the morphogenetic potential of their actin cytoskeletons. Keratinocytes demonstrated the most complex actin cytoskeleton organization as compared to non-keratinocyte epithelial cells. At the same time the process of epithelial-mesenchymal transition partially restored the ability of non-keratinocyte epithelial cells to assemble a complex actomyosin network. Of note, the degree of development of such networks also correlated with EMT score (computed using measurement of multiple parameters related to gene expression). Thus, analysis of the cytoskeleton organization in a well-defined microenvironment may provide an additional tool for monitoring the diversity of cancer cell populations.

## Materials and methods

### Cell culture, treatment and transfection

All cells were maintained in a humidified incubator set at 37°C and atmospheric 5% CO_2_. Human foreskin fibroblasts (HFFs) were cultured and transfected as previously described ^15^. NBT-II, MCF-7, HaCaT, CaCo-2, MDCK, SKOV3 and OVCA429 were cultured in Dulbecco’s Modified Eagle’s Medium (DMEM, Invitrogen) high glucose supplemented with 10% Heat Inactivated Fetal Bovine Serum (HI-FBS, Invitrogen), 1mM sodium pyruvate (Invitrogen) and 0.1% penicillin and streptomycin (P/S, Invitrogen). MCF-10A were cultured in DMEM:Nutrient Mixture F12 (DMEM/F12, Invitrogen) supplemented with 5% horse serum (Invitrogen), 20ng/ml Epidermal Growth Factor (EGF, Peprotech), 0.5mg/ml Hydrocortisone (Sigma), 100ng/ml Cholera Toxin (Sigma), 10mg/ml Insulin (Sigma) and 0.1%P/S. Murine keratinocytes (Wild Type and Knockout, kind gifts from T. Magin Group, University of Leipzig, Germany) were cultured in low Ca^2+^ DMEM/ Ham’s F12 (Merck) supplemented with 10% chelex-treated HI-FBS, 10ng/ml EGF, 0.5mg/ml hydrocortisone, 10^-10^M Cholera Toxin, 2.5mg/ml insulin, 0.18mM adenine (Sigma), 1mM Glutamax (Invitrogen), 1mM sodium pyruvate and 0.1% P/S. HEYA8 and PEO1 were culture in Roswell Park Memorial Institute-1640 media supplemented with 10% HI-FBS and 0.1% P/S.

To induce EMT in epithelial cells, NBT-II were treated in culture media supplemented with 100ng/ml EGF while HaCaT were treated in culture media supplemented with 20ng/ml TGF-β1 and 100ng/ml EGF for one day and refreshed with 10ng/ml TGF-β1 and 100ng/ml EGF daily for three more days before overnight imaging. To induce chirality in keratinocytes, HaCaT were treated with low doses 20-200nM of latrunculin A (Sigma).

Transient transfection of DNA plasmids into cells was done by electroporation (Neon transfection system, Life Technologies) following manufacturer’s instrcutions. Electroporation conditions for NBT-II, MCF-10A and HaCaT were 2 pulses of 1200V for 20ms, while MCF-7 required 2 pulses of 1250V. Plasmids encoding the following fluorescence fusion proteins were used: Lifeact-GFP (R. Wedlich-Soldner Group, Max Planck Institute of Biochemistry, Martinsried, Germany), RFP-F-Tractin (Zaidel-Bar Group, Mechanobiology Institute, Singapore), mRuby-Lifeact and mEmerald-Keratin14 (Michael W. Davidson Group, Florida State University, Tallahassee, FL, USA).

### Protein micropatterning of substrates

Cells were seeded on substrates containing circular adhesive islands of varying area (500, 750, 1000, 1500, 2000 and 2500μm^2^), or, circular islands with fixed areas (700, 1200 or 1800 μm^2^). Adhesive circular islands were fabricated by using a PDMS stamp in either micro-contact printing as described previously ^15^, or, by a slightly modified version of stencil patterning ^44^. In stencil patterning, PDMS stamps were first inverted and placed onto ibidi hydrophobic uncoated 35mm μ-dishes (ibidi GmbH). NOA-73 (Norland Inc.) was deposited along an edge of the stamp and allowed to fill in the gaps between the PDMS stamp and dish by capillary action. The NOA stencil was cured under ultraviolet illumination for ~15s. After peeling off the PDMS stamp, the stencil and dish were incubated with fibronectin (Calbiochem, Merck Millipore) or collagen I (BD Bioscience) at a concentration of 50μg/ml in PBS or acetic acid respectively at 4°C overnight. Unadsorbed protein was rinsed off, the NOA stencil removed and the dish was then passivated with 0.2% pluronic acid in water for 10mins at 37°C. Finally dishes were rinsed in PBS thrice before epithelial cells were seeded at a density of 6 or 7×10^4^ cells/ml, while fibroblasts were seeded at 5×10^4^ cells/ml.

### Immunofluoroscence and immunoblotting

Cells were fixed with 4% paraformaldehyde in PBS for 10mins, followed by three PBS washes. Cells fixed with paraformaldehyde were permeabilized with 0.5% Triton-X-100 and subsequently quenched with 0.1M glycine in PBS for 10mins each. After PBS washes, blocking was done with 2% BSA in PBS for 1hr at Room Temperature (RT) prior to overnight primary antibody incubation at 4°C with mouse anti-paxillin (1:100, BD Bioscience) or anti-β-actin (1:200, Sigma) in 2% BSA in PBS. Fixed cells were washed with PBS three times and then incubated with an appropriate Alexa Fluor-conjugated mouse secondary antibody (1:250 dilution, Thermo Fisher Scientific) in 2% BSA in PBS for 1hr at RT. F-actin staining was done using Alexa Fluor 488 (Thermo Fisher Scientific) or TRITC (Sigma) conjugated Phalloidin at a dilution of 1:500 while Keratin5 staining was done using anti-cytokeratin5 conjugated to Alexa Fluor 647 at a dilution of 1:100 (Abcam), incubated overnight at 4°C or 1hr at RT.

Growth factor treated cells were lysed with RIPA buffer (Sigma) and extracted proteins were separated by 4–20% gradient SDS-polyacrylamide gel electrophoresis (Thermo Fisher Scientific) and transferred to a 0.2μm PVDF membrane (Bio-Rad) at 100V for 1.5hrs. Membranes were blocked with 5% non-fat milk (Bio-Rad) in TBS for 1hr at RT before incubation with the appropriate primary antibody: mouse anti-GAPDH (Santa Cruz) at a dilution of 1:3,000, mouse anti-E-cadherin (BD Transduction) at a dilution of 1:5,000, rabbit anti-slug (Cell Signaling Technology, CST) at a dilution of 1:250, rabbit anti-vimentin (CST) at a dilution of 1:250 in 5% non-fat milk in PBS at 4°C overnight. After washes in TBST, the membrane was incubated with an appropriate HRP-conjugated secondary antibody (Bio-Rad) at a dilution of 1:2000 in 2.5% non-fat milk in PBS. The membrane was then processed for ECL detection (Bio-Rad) and chemiluminescence was detected in the Bio-Rad ChemiDoc Imaging System.

### Imaging, processing and statistical analysis

For live imaging, cells cultured in phenol red containing DMEM were switched to Leibovitz’s L-15 (supplemented with 10% FBS and 1mM sodium pyruvate), while MCF-10A was imaged in normal culture media buffered by supplementing a final concentration of 20mM HEPES (Invitrogen) and 0.369% sodium bicarbonate (Invitrogen). Time-lapse imaging was done at imaging rates of 2-10mins/frame with z-stacks (10-15μm) of step size 0.35μm, on a spinning-disc confocal microscope (PerkinElmer Ultraview VoX) on an inverted microscope (Olympus IX81) using 60x (UPlanSApo, NA 1.2, water) or 100x (UPlanSApo, NA 1.4, oil) objectives. Image acquisition was set up in Volocity software and taken by an EMCCD camera (Hamamatsu C9100-13). Brightfield images where taken on the EVOS FL Imaging system (Thermo Fisher Scientific) using the 4x objective.

Maximum intensity projection through the z-stack of time-lapse images was done in Volocity before exporting as TIFF files for further processing in ImageJ (National Institutes of Health, NIH). Image processing functions applied to whole images included: Bleach correction, Gaussian Blur Filter, Sharpen, Subtract Background. For measurement of cell spread area and total focal adhesion area, segmentation and particle detection of single plane phalloidin and paxillin stained images was done in ImageJ using the Adaptive 3D Threshold and Analyze Particles functions respectively.

Graph plotting and statistical analysis were performed in Prism. Analyses of significant differences were carried out using two-tailed Mann-Whitney test or one-way ANOVA with Tukey’s multiple comparison test for more than two groups. Statistical methods and sample size were specified in figure legends and differences were accepted as significant if *p*<0.05. No statistical method was used to predetermine sample size.

## Acknowledgements

We are grateful to the Magin Group at the University of Leipzig as well as the Low, Zaidel-Bar and Ladoux Groups at the Mechanobiology Institute (MBI) for graciously providing us with epithelial cell lines. We thank the Kanchanawong Group at MBI for managing and distributing the M. Davidson groups’ plasmid inventory. We appreciate the microscopy core and wet lab core facility at MBI for technical help and providing equipment. This research is supported by the National Research Foundation, Prime Minister’s Office, Singapore and the Ministry of Education under the Research Centres of Excellence programme (R-714-006-006-271). S.J. was funded by the Mechanobiology Institute, National University of Singapore through a graduate scholarship.

## Supplemental Figure Legends

**Figure S1:**
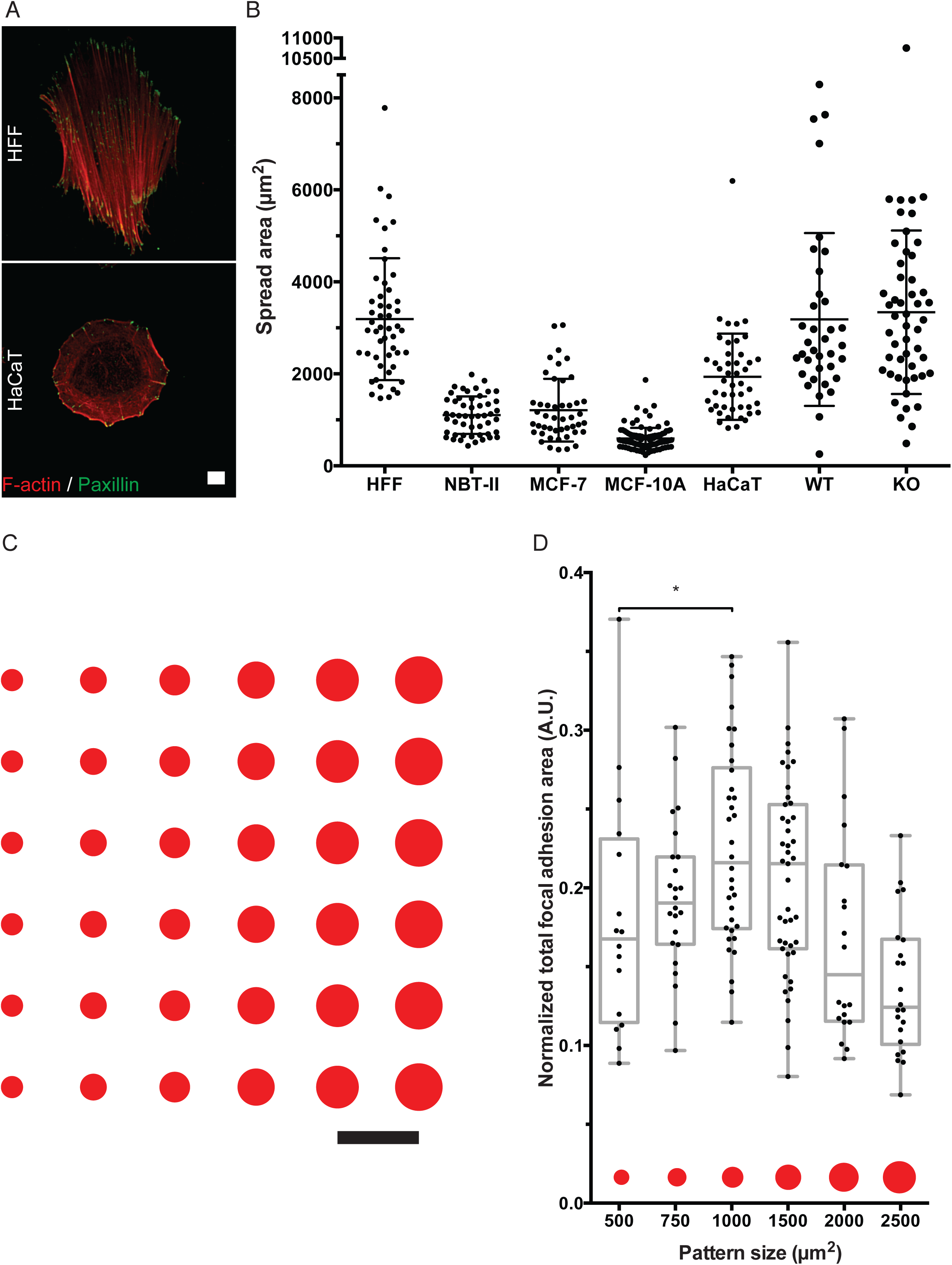
Cell morphology and spread area variation between epithelial cell lines and fibroblasts. **A)** Representative images showing focal adhesions (paxillin immunostaining) and F-actin (phalloidin staining) distribution in HFF and HaCaT fixed 24 hours after seeding on planar fibronectin coated surfaces. Scale bar, 10μm. **B)** Scatter dot plot with Mean ± Standard Deviation showing cell spread area of different cell lines on planar protein coated surfaces. HFF, NBT-II and HaCaT were fixed and stained with phalloidin 24hrs after seeding on planar fibronectin coated surfaces. Wild Type (WT) and Knockout (KO) murine keratinocytes were fixed and stained with phalloidin 22hrs after seeding on planar collagen coated surfaces. MCF-7 and MCF-10A were fixed and stained with phalloidin 18hrs after seeding on planar collagen coated surfaces. **C)** Schematic showing pattern consisting of six different sizes of circular islands that was used for protein patterning. Scale bar, 50μm. **D)** Box and whiskers plot showing the total focal adhesion area in cells imaged under conditions of Figure 1C normalized by cell projected area.

**Figure S2:**
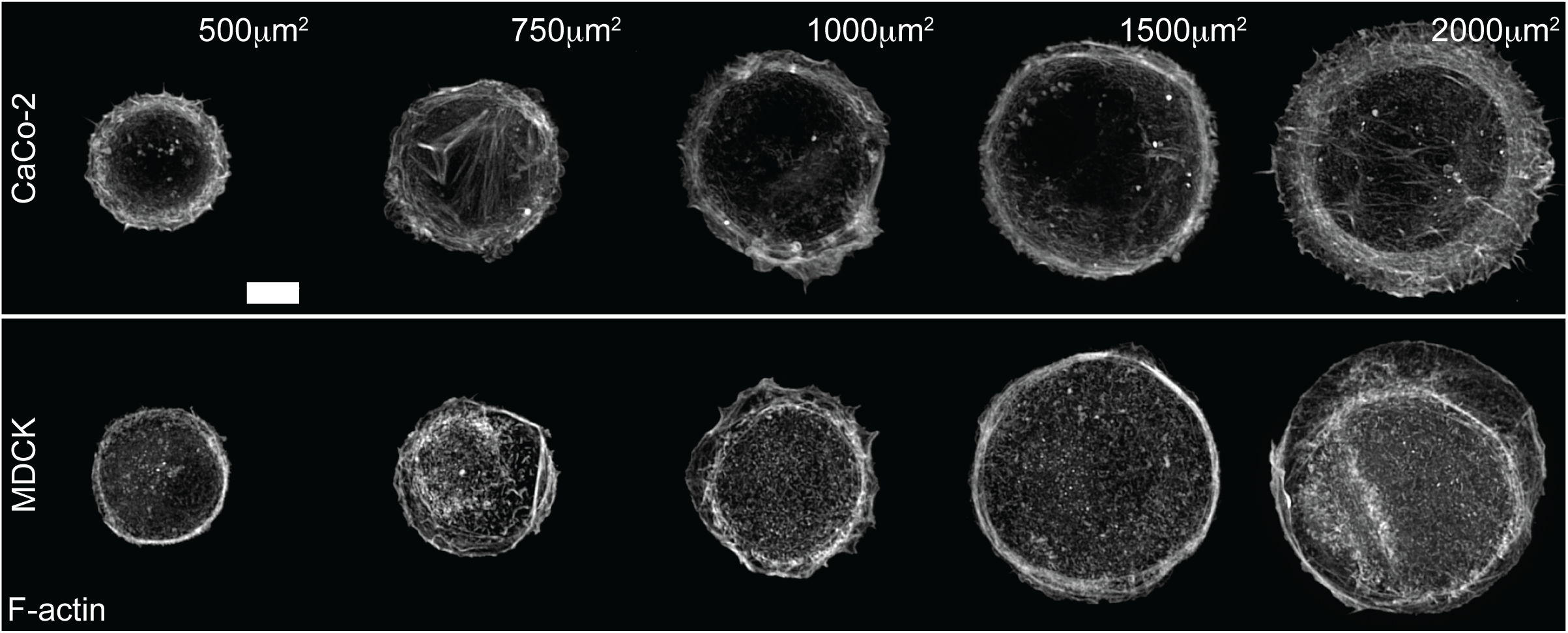
Actin cytoskeleton development in non-keratinocyte epithelial cell lines. Representative images showing phalloidin staining in Caco-2 and MDCK fixed 20-24hrs after seeding on collagen coated islands of different sizes. Scale bar, 10μm.

**Figure S3:**
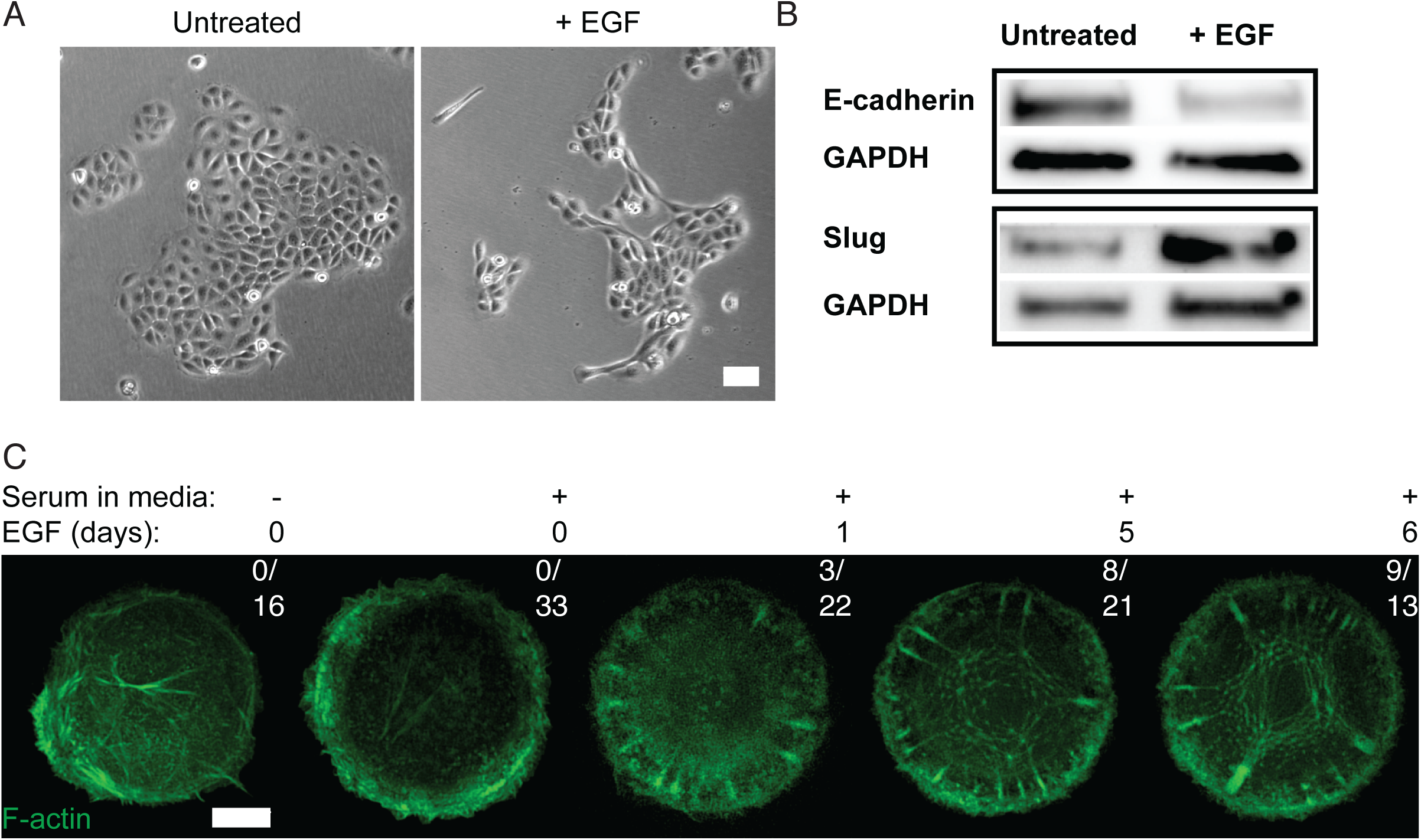
Actin cytoskeleton development during epithelial-mesenchymal transition in rat bladder carcinoma cells. **A)** Representative brightfield images of NBT-II in standard culture after cells have not been previously treated with anything (Untreated, left panel) or have been previously treated with 100ng/ml EGF for 3 days (+EGF, right panel). Scale bar, 100μm. **B)** Western blot images showing protein expression levels of E-cadherin, slug and GAPDH in total protein lysates taken from NBT-II cells without (Untreated) or with (+EGF) 100ng/ml EGF treatment for 3 days. **C)** Representative frames of NBT-II in serum free or serum containing media with or without 100ng/ml EGF pre-treatment (in days) expressing Lifeact-GFP seeded on 1200μm^2^ fibronectin coated islands and imaged overnight. Fractions of cells able to develop radial fibres (over total number of imaged cells) are shown in the top right hand side of each frame representing a treatment condition. Scale bar, 10μm.

**Figure S4:**
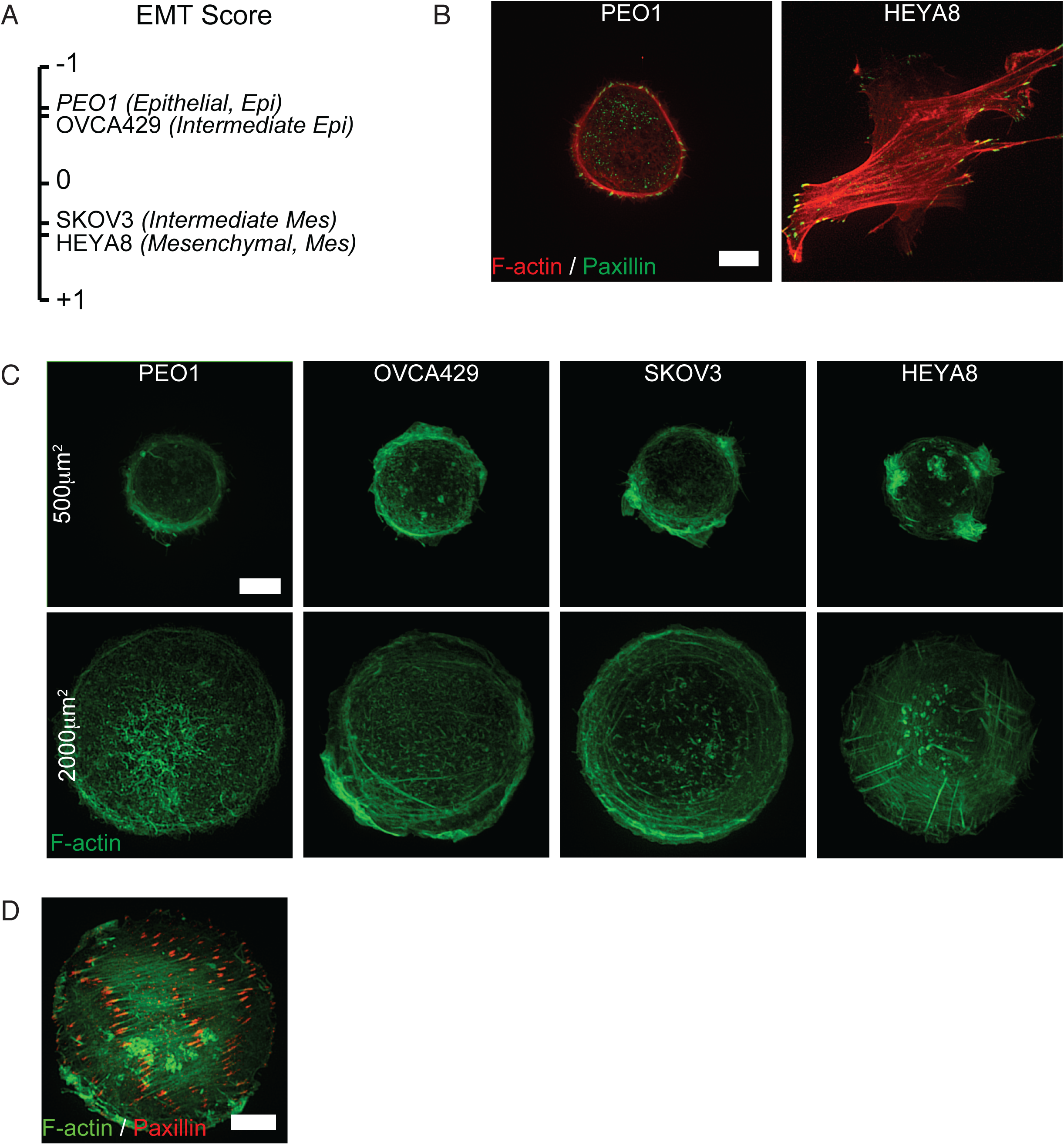
Actin cytoskeleton self-organization in ovarian cancer cell lines. **A)** A selection of ovarian cancer cell lines (PEO1, OVCA429, SKOV3 and HEYA8) distributed along a scale corresponding to their EMT score (from most epithelial, -1 to most mesenchymal, +1). **B)** Representative images showing focal adhesion (paxillin immunostaining) and F-actin (phalloidin staining) distribution in epithelial (PEO1) and mesenchymal (HEYA8) ovarian cancer cells fixed 16 hours after seeding on planar fibronectin coated substrates. Scale bar, 10μm. **C)** Representative images of ovarian cancer cells with different EMT phenotypes (PEO1, OVCA429, SKOV3 and HEYA8) fixed and stained with phalloidin 16hrs after seeding on fibronectin coated islands of different sizes. Scale bars, 10μm. **D)** Representative image showing focal adhesion (paxillin immunostaining) and F-actin (phalloidin staining) distribution in a HEYA8 cell imaged under conditions detailed in **C** that shows the linear actin pattern. Scale bar, 10μm.

**Figure S5:**
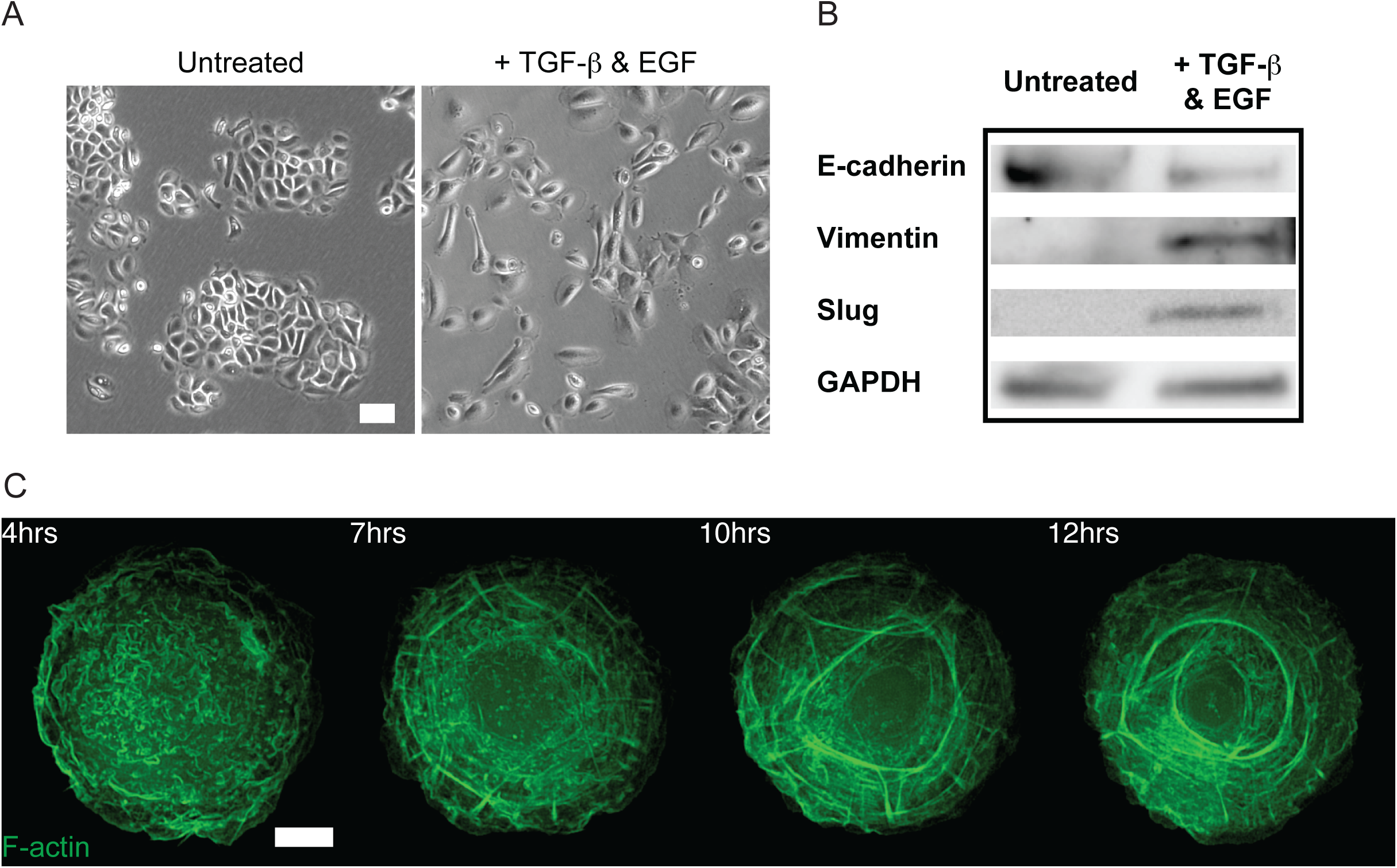
Epithelial-mesenchymal transition in human keratinocytes. **A)** Representative brightfield images of HaCaT in standard culture after cells have not been previously treated with anything (left panel) or have been previously treated with EGF and TGF-β for 3 days (as described in Materials and Methods, right panel). Scale bar, 100μm. **B)** Western blot images showing protein expression levels of E-cadherin, vimentin, slug and GAPDH in total protein lysates taken from HaCaT cells without (Untreated) or with (+TGF-β & EGF) growth factor treatment for 5 days. **C)** Timlapse series of actin pattern development in HaCaT cells replated on 1800μm^2^ fibronectin coated islands of different sizes after 4 days of growth factor treatment. Cells were transfected with Lifeact-GFP one day before replating and imaging overnight. Time in hours after seeding is indicated in upper left corner of each frame. Scale bar, 10μm.

## Abbreviations List

EMT: epithelial-mesenchymal transition
EGF: epidermal growth factor
WT: Wild Type
KO: Knockout
RF: radial fibre
TF: transverse fibre

